# EVOH Hollow Fiber Encapsulation of Individual *Caenorhabditis elegans* for Transmission Electron Microscopy

**DOI:** 10.1101/2025.10.08.681240

**Authors:** Willisa Liou, David M. Belnap

## Abstract

We present a streamlined protocol for encapsulating individual *Caenorhabditis elegans* specimens for thin-section transmission electron microscopy (TEM) using ethylene vinyl alcohol (EVOH) copolymer hollow fibers. These transparent fibers, featuring a 175 μm inner diameter, 25 μm wall thickness, and an approximate molecular weight cutoff of 30 kDa, possess both hydrophilic and hydrophobic properties. This amphiphilic nature helps maintain a potent lumen structure even in dry, open air, enabling precise handling and orientation of small specimens prior to resin polymerization and eliminating the need for remounting. The protocol includes specimen transfer using wire loops, microwave-assisted chemical pre-fixation, dissection on a soft silicone substrate with micro-scissors, and manipulation of hollow fibers using insect pins. This approach preserves the identity of individual specimens throughout processing. During ultramicrotomy, the fiber serves as a visual guide, enhancing alignment and reproducibility of sectioning. As proof of principle, we demonstrate well-preserved ultrastructural features in chemically fixed, microwave-assisted resin-embedded *C. elegans* specimens. This method offers a robust and reproducible alternative to conventional gelatinous media encapsulation, particularly suited for TEM analysis of scarce or small biological samples.

## INTRODUCTION

The free-living soil nematode *Caenorhabditis elegans* is a well-established model organism in biological research. Owing to its small size—approximately 1 mm in length and 50 μm in diameter in adults (Maguire et al., 2011; Moore et al., 2013)—transmission electron microscopy (TEM) has been extensively employed to investigate its internal architecture. The depth of TEM-based studies has led to the development of a publicly accessible ultrastructural atlas of the wild-type animal (WormAtlas, 2025), which includes a complete map of its neuronal wiring (Cook 2019; White et al., 1986). EM continues to be an indispensable tool for characterizing structural abnormalities, particularly in neuronal mutants (Hsu et al., 2014; Zheng et al., 2017).

Preparation of biological samples as thin sections for TEM involves a complex, multi-step process with successive chemical treatments. When working with minute specimens such as *C. elegans*, direct handling with forceps can easily damage their delicate anatomical structures. The most common approach to increasing specimen size has been encapsulation in melted gelatinous media. For resin-based preparations, agar (Ward et al., 1975) or low-melting-point agarose (Hall et al., 2012; Wood and Klomparens, 1993) are preferred because they remain transparent after osmication. In contrast, gelatin is typically used for cryosection preparations (Sato et al., 2005). Once cooled, the solidified matrix encapsulating the specimen is trimmed to a manageable size for further processing. These methods aim to prevent the loss of valuable specimens during solution changes and transfers. However, these gelatinous procedures are themselves tedious (Sigmond et al., 2008) as they require sequential steps of heating, embedding, cooling, and trimming prior to sample dehydration. Furthermore, for non-resin-embedding preparations intended for cryosectioning (Tokuyasu method, Tokuyasu 1973), manually trimming the semi-fluid encapsulated sample to achieve precise orientation prior to freezing is particularly challenging (Nicolle et al., 2015; Sato et al., 2005).

In this study, we introduce an alternative encapsulation method using hollow fibers to prepare *C. elegans* specimens for thin-section TEM examinations. Hollow fibers, typically housed within a hemodialyzer (commonly known as an artificial kidney), function as dialysis membranes in the form of capillary tubing. These semi-permeable membranes have long been used in molecular separation techniques.

Nordbring-Hertz et al. (1983; 1984) innovatively employed cellulose dialysis membrane strips (1–2 cm²) as both a growth matrix on agar surfaces and a transport medium for nematode-trapping fungi and their nematode prey. This approach enabled the transfer of intact fungal-nematode complexes for electron microscopic studies without physically disturbing or damaging the nematodes, simply by handling the membrane itself.

Hohenberg et al. (1994) pioneered the use of cellulose capillary tubes, 200 μm in diameter, to confine living nematodes specifically for cryoimmobilization. The encapsulated animals were subsequently processed via freeze-substitution, a chemical exchange technique that yielded resin blocks suitable for TEM analysis. While capillary action facilitated the uptake of liquid-suspended specimens into the tubes, the method was inherently complex due to its aim of achieving instantaneous cryoimmobilization. Loading worms into the narrow tubes required considerable dexterity and the use of specialized devices (Claeys et al., 2004; Müller-Reichert et al., 2003). Alternative approaches to cryoimmobilization were later developed (Leunissen and Yi, 2009; Müller-Reichert et al., 2003), but all shared the challenge of handling live specimens under stringent conditions.

Rather than aiming for instantaneous cryoimmobilization techniques such as plunge or high-pressure freezing, we adopted the concept of encasing minute specimens in small tubes for downstream electron microscopy processing. Specifically, this report focuses on the technical implementation of worm encapsulation using non-cellulose-based hollow fibers extracted from a hemodialyzer. This approach offers a practical and versatile alternative to gelatinous encapsulation methods and is particularly suitable for scarce or small biological samples. Hydrated specimens encapsulated in hollow fibers can be readily cryosectioned using Tokuyasu-style protocols (Tokuyasu, 1973), bypassing resin embedding and enabling efficient preparation for ultrastructural immunolocalization at the electron microscopy level.

## MATERIALS AND METHODS

The workflow for enclosing *C. elegans* in EVOH hollow fiber for TEM analysis is summarized in Figure 1 and demonstrated in Videos 1–6.

**Figure 1.**
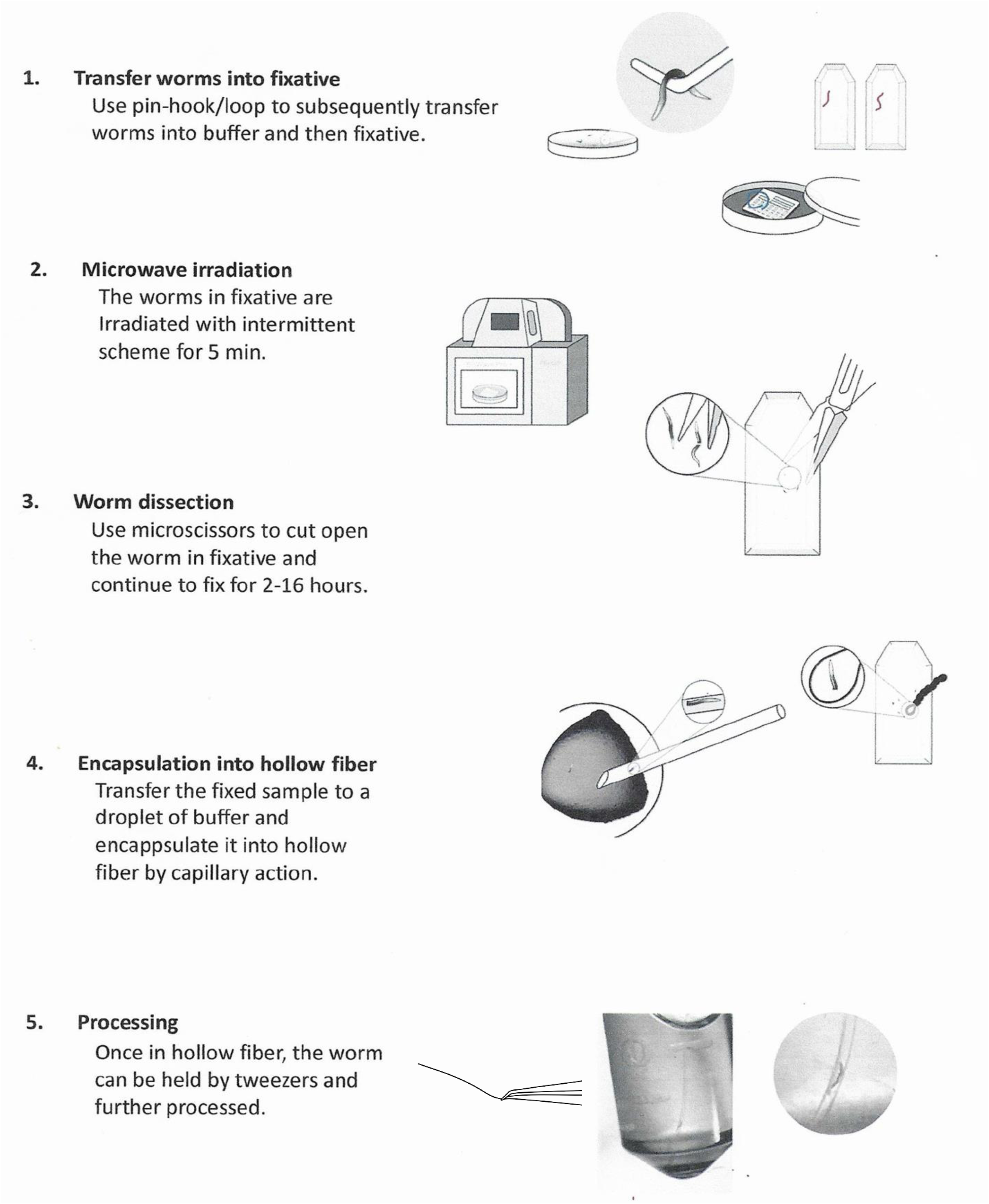
Essential steps of worm encapsulation

### Step1. Worm transfer

#### *C. elegans* Cultivation and Transfer Using Pin-Hook Technique (video 1)

Wild-type *Caenorhabditis elegans* (N2 strain), obtained from the Caenorhabditis Genetics Center (CGC), were maintained on Nematode Growth Medium (NGM) plates seeded with Escherichia coli OP50 as a food source (Brenner, 1974). Upon depletion of the bacterial lawn, worms were transferred to fresh plates using a sterilized pin-hook (see below) disinfected with 70% ethanol. All manipulations were performed under a stereo dissecting microscope equipped with transmitted illumination from below.

#### Fabrication and Use of Insect Pins and Wire Loop (video 2 and 3)

Stainless steel insect pins (Minutiens, Austerlitz Insect Pins®; ∼12 mm length, 0.1 mm shaft diameter, 0.0125 mm tip diameter) were used as worm picks and for hollow fiber handling. The blunt ends were annealed to P10 pipette tips for ease of manipulation; pin tips were shaped into hooks, loops, or clips depending on the task (Fig. 2A, left and middle panels). To facilitate individual worm tracking during liquid transfers, a wire loop was employed instead of standard pipetting techniques. This approach leveraged the surface tension property that allows a small droplet of fluid to remain suspended in air. The loop (Fig. 2A, right panel) was constructed by twisting a platinum wire (ϕ 0.008”; Ted Pella Inc., #23-2) around a hypodermic needle to form an inner diameter of approximately 0.5 mm. This configuration enabled the capture and transfer of individual swimming worms or fixed worm fragments within the droplet

**Figure 2:**
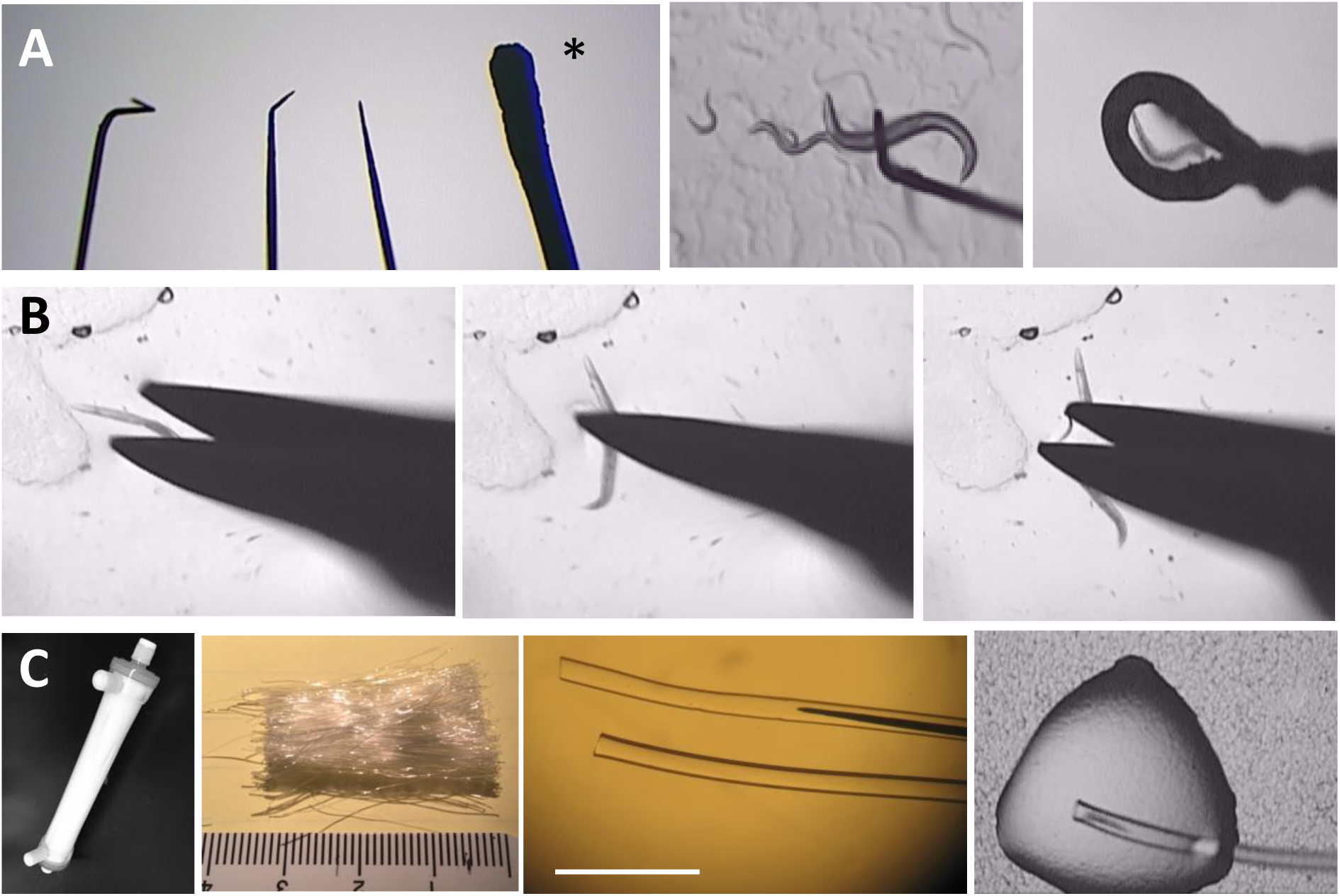
**Tools and procedures** A: *insect pins and platinum loop* (left panel) pins are bent to various shapes; a platinum worm pick is included at the right (*) as a comparison of dimensions; (middle panel) a pin hook is placed betweem a worm and agar plate ready for pick- up; (right panel) a worm is suspended in a droplet in a loop. B: *microscissors:* the blades are approaching a worm → a snap is being made → the worm is divided. C: *hollow fiber*: (from left to right): a hemodialyzer (33x5 cm); a bunch of hollow fibers retrieved from a hemodialyzer and segmented into 3 cm length; hollow fiber with and without an insect pin inserted; a half-worm specimen being pulled into the fiber by capillary action. bar: 5 mm

#### Silicone Platform for Specimen Manipulation

A transparent, flat silicone embedding mold (Ted Pella, Inc., #10504)—typically used for resin embedding—served as the platform for worm manipulation. The mold features multiple wells in three distinct sizes, allowing buffer and fixative solutions to be kept nearby yet separated. The silicone rubber material is soft and pliable, enabling direct dissection of worms with scissors on its surface. Additionally, due to the hydrophobic nature of silicone, small volumes of buffer form droplets with high contact angles, which facilitates encapsulation procedures.

#### Transfer and Preparation for Fixation

Single young adult worms were picked and rinsed in 250 μl of Life Cell Imaging Buffer (ThermoFisher #A1429IDJ), then transferred with a wire loop to 250 μl of a half-strength fixative solution. This diluted fixative was prepared as a 1:1 mixture of the full-strength fixative (2% glutaraldehyde, 0.64% paraformaldehyde in buffer: 21 mM PIPES, 8 mM HEPES, 56 mM NaCl, 1 mM KCl, 0.72 mM CaCl₂, 0.4 mM MgCl₂) and distilled water.

### Step 2. Pre-fixation

#### Microwave-Assisted Pre-Fixation

Although *C. elegans* is small, its cuticle presents a significant barrier to chemical fixatives. To facilitate rapid penetration into internal tissues and prevent autolysis, the animals must be incised. However, dissection of live worms frequently results in rupture of the intestine and gonads. To mitigate this, worms were pre-fixed by microwave irradiation for 10 minutes while immersed in half-strength fixative (see above, Table 1, Step 1) prior to dissection. For safety, the silicone mold containing the worms submerged in fixative was covered with a glass Petri dish during irradiation, and the microwave was operated with active air exhaust.

**TABLE 1.** Microwave assisted chemical fixation and sample processing protocol

| step |  | solution | Irradiation Power (watt) | Irradiation time | restriction temperature | Probe location | repeat |
| --- | --- | --- | --- | --- | --- | --- | --- |
| Chemical fixation | 1 | Half-strength fixative<sup>‡</sup> | 125 | 1.5 min* | 37°C | reservoir | 5x |
|  | 2<sup>*</sup> | Full-strength fixative | Fixed at room temperature for 2-16 hours, no irradiation. |  |  |  |  |
| Processing in microcentrifuge tubes | 3 | buffer | 125 | 1 min | 37°C | reservoir | 1x |
|  | 4 | 2% osmium tetroxide in buffer | 125 | 5 min | 37°C | reservoir | 1x |
|  | 5 | distilled water | 125 | 1 min | 37°C | reservoir | 1x |
|  | 6 | 4% aqueous uranyl acetate | 125 | 5 min | 37°C | reservoir | 1x |
|  | 7 | 50% alcohol | 250 | 1 min | 37°C | blank | 1x |
|  | 8 | 70% alcohol | 250 | 1 min | 37°C | blank | 1x |
|  | 9 | 95% alcohol | 250 | 1 min | 37°C | blank | 1x |
|  | 10 | 100 % alcohol | 250 | 1 min | 37°C | blank | 5x |
|  | 11 | 100 % acetone | 250 | 1 min | 37°C | blank | 5x |
|  | 12 | 1:1 resin: actone | 250 | 15 min | 45°C | blank | 1x |
|  | 13 | 3:1 resin:acetone | 250 | 15 min | 37°C | blank | 1x |
|  | 14<sup>¶</sup> | resin | 250 | 15 min | 37°C | blank | 3x |

<sup>†</sup> PELCO BioWave<sup>®</sup> Pro 36500 Laboratory Microwave System (Ted Pella, INC). All irradiation steps are performed inside the oven on the Pelco Coldspot device.
<sup>‡</sup> Due to the small volume of the fixative, it is diluted with an equal amount of distilled water to counteract evaporation.
\* Intermittent irradiation scheme per cycle: 10 sec on 20 sec off.
<sup>×</sup> the worm is cut open when step 1 is finished.
<sup>¶</sup> under vacuum.

### Step 3 Worm Dissection

#### Worm Dissection Using Micro Scissors on Silicone Surface (video 4)

Following initial chemical fixation, worms became motionless and settled at the bottom of the silicone cavity. Dissection was performed using spring-loaded micro scissors (Vannas H- 4240, Albert Heiss GmbH & Co. KG). The scissors were lowered vertically into the well, targeting the worm. Owing to hydrophobic interactions between the worm cuticle and the silicone surface, the specimen remained stationary despite fluid disturbance caused by the approaching scissors. With the blades positioned around the worm, the tips of the scissors were gently pressed into the silicone. A snip was made at the mid-body or near the ends (Fig. 2B). The dissected worm then remained in the same well for 2–16 hours at room temperature while the fixative was replenished to full-strength (Table 1, Step 2)

#### Preparation of Hollow Fibers for Encapsulation

Hollow fibers (inner diameter: 175 μm; wall thickness: 25 μm) were extracted from a KF- 201 hemodialyzer (Fig. 2C; Kawasumi Laboratories, Inc.) by cutting open the unit and separating the fiber bundle. The fibers, made of ethylene vinyl alcohol (EVOH) copolymer with ∼30 kDa molecular weight cut-off, were cut into ∼3 cm segments. An unbent insect pin (100 μm diameter), annealed to a pipette tip, was inserted into the fiber lumen for manipulation. This setup allowed precise control during specimen loading and transfer.

### Step 4 Encapulation

#### Encapsulation Procedure (video 5)

Approximately 3 μl of rinsing buffer was placed on the silicone mold. Using a wire loop and an eyelash tool, a piece of fixed specimen was deposited onto the buffer droplet. A hollow fiber, held on an insect pin, was brought into contact with the droplet, allowing fluid and the worm to be drawn into the fiber via capillary action (Fig. 2C, right-most panel). Once loaded, the fiber was promptly submerged in a nearby larger well containing ample buffer to prevent air bubble entrapment. The insect pin was gradually withdrawn by leaning it against the edge of the silicone well, which ensured the buffer fully filled the fiber and prevented air bubble formation. The loaded fiber was left submerged for 3–5 minutes to facilitate thorough rinsing of residual fixative and equilibration in the aqueous environment.

During this time, the fiber was trimmed and sealed at both ends—either by compression with forceps or by heating with a platinum wire—to prevent specimen escape. If the worm was positioned centrally within the fiber, sealing was deemed unnecessary because the fiber’s length exceeded the specimen’s by more than 20-fold, effectively preventing escape. No differences in chemical exchange were detected between sealed/short and unsealed/long fibers.

The encapsulation procedure was completed within ten minutes. The encapsulated specimen, hereafter referred to as a capsulate, was handled with tweezers and processed according to standard tissue specimen protocols.

### Step 5 Processing

#### Microwave-Assisted sample processing

Capsulates were transferred to polypropylene microcentrifuge tubes and processed using a microwave-assisted protocol (Table 1, Steps 3–14). Solutions were exchanged by gently pipetting from the tube while leaving the hollow fiber undisturbed. When the Embed 812 resin (EMS #14120) became sufficiently viscous during the final stages of infiltration, it was no longer possible to aspirate the resin efficiently. At this point, capsulates were grasped with forceps and moved to a fresh tube containing new resin to ensure complete infiltration. If a specimen exited the hollow fiber during processing, solution exchanges were continued through step 12, after which the specimen was re-encapsulated in a 1:1 resin-acetone mixture (Video 6).

### Step 6. Embedding, sectioning and imaging

Following complete infiltration of the resin, the capsulate was transferred to a horizontal embedding mold. Excess fiber length was trimmed to achieve the desired orientation of the specimen, and the sample was cured overnight at 65 °C. Polymerized samples were sectioned using a Leica EM UC6 ultramicrotome. Semithin sections (∼0.5 μm) were collected on glass slides and stained with 1% toluidine blue in 1% boric acid. Light microscopy images were acquired using an Axio Scan.Z1 system (Zeiss). Ultrathin sections (∼40–60 nm) were collected on Pioloform/Formvar-coated slot grids and post-stained with aqueous uranyl acetate followed by Reynold’s lead citrate. Transmission electron microscopy (TEM) images were obtained using a JEOL JEM-1400 microscope operated at 120 kV and equipped with a Gatan Orius digital camera. For electron tomography, 80 nm sections were mounted in a high-tilt specimen holder (EM 21311HTR). Serial tilt images (±60°, at 2° increments) of the region of interest were acquired using SerialEM (Boulder Laboratory for 3-D Fine Structure). Three-dimensional reconstructions of tilt series were computed using IMOD software (Kremer et al., 1996).

## RESULTS

### Accelerated Processing and Ultramicrotomy

Hollow fiber encapsulation, when combined with microwave-assisted pre-fixation and processing, significantly reduces manual handling time to under two hours. Integrating the hollow fiber during the embedding step enables convenient adjustment of sample orientation prior to resin polymerization (Fig. 3A), thereby eliminating the need for re-embedding after curing. During ultramicrotomy, the fiber also serves as a visual guide for precise trimming. As shown in Figs. 3B–F, the worm remains suspended within the fiber, elevated above the mold base, creating a buffer zone equivalent to at least the thickness of the fiber wall (≥25 μm). This clearance facilitates pyramid trimming, which is essential for producing ultrathin sections.

**Figure 3:**
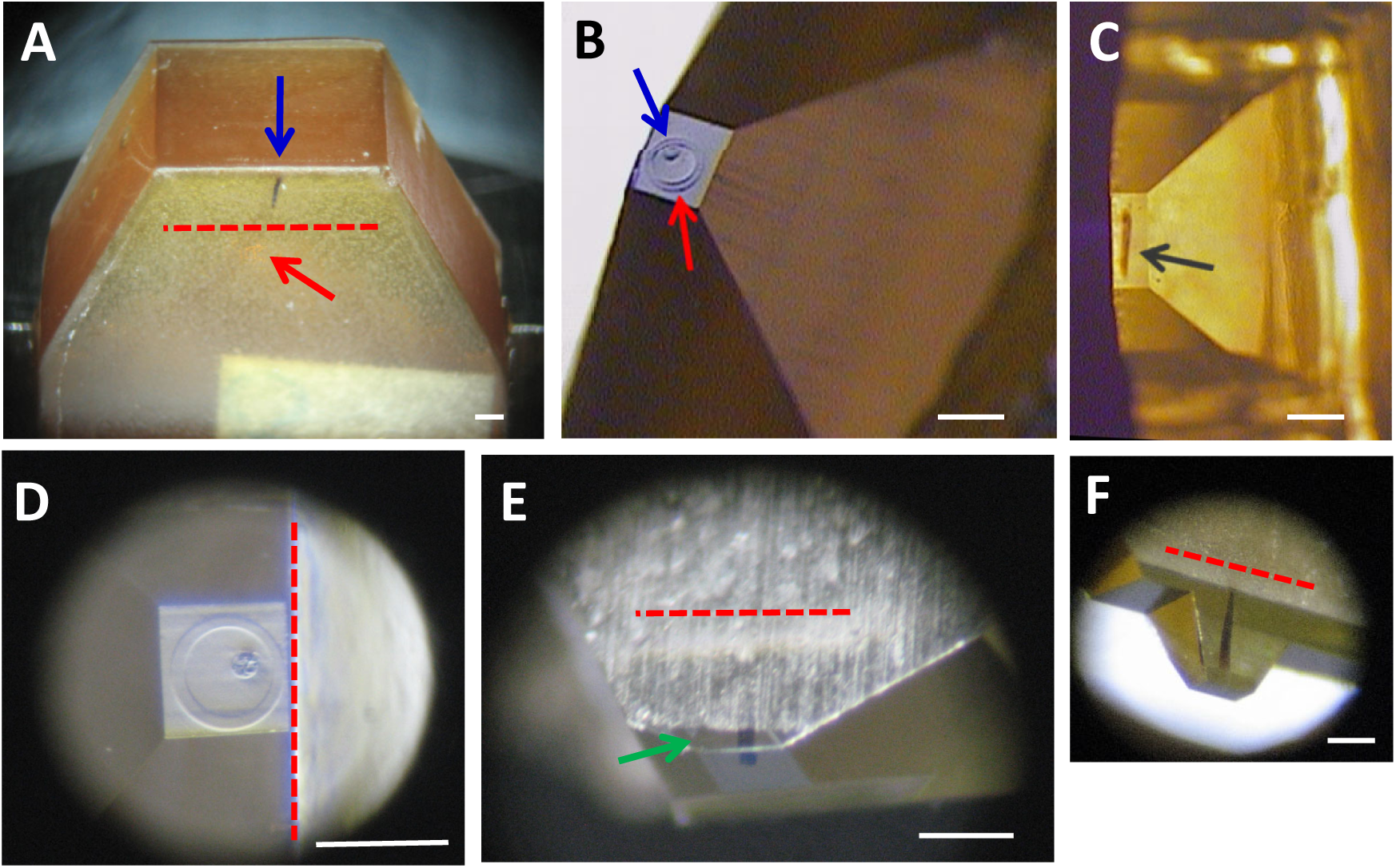
Ultramicrotomy is expedited. The worm specimen is well-oriented, and block trimming on all four sides is possible without re-mounting the specimen. A: an untrimmmed resin block showing that the entire capsulate has sunken into the bottom side of the block. C- F: trimmed block. Note that the rugged bottom surface can be trimmed smoothly without jeopardizing the worm. Red-dashed line denotes the bottom surface of the block. (red arrow = EVOH fiber; blue arrow = worm; black arrow = longitudinally sectioned worm. green arrow = slightly trimmed bottom surface.) bars: 250 μm

The EVOH proved to be a resilient encapsulation medium. The fiber maintained its structural integrity throughout microwave irradiation, chemical infiltration, and heat exposure. In Fig. 4A, a semithin (0.5 μm) section stained with toluidine blue reveals that the fiber retained a distended, rounded contour, consistent with equilibration of diffusible substances across the membrane. The worm was eccentrically positioned within the lumen, occupying approximately 10% of the enclosed area. This spatial configuration allows for trimming of more than 50 μm from all four sides of the resin block without cutting into the specimen, thereby facilitating serial sectioning. Figure 4 presents corresponding light and electron micrographs at increasing magnifications, demonstrating the continuity and high quality of the resulting sections.

**Figure 4:**
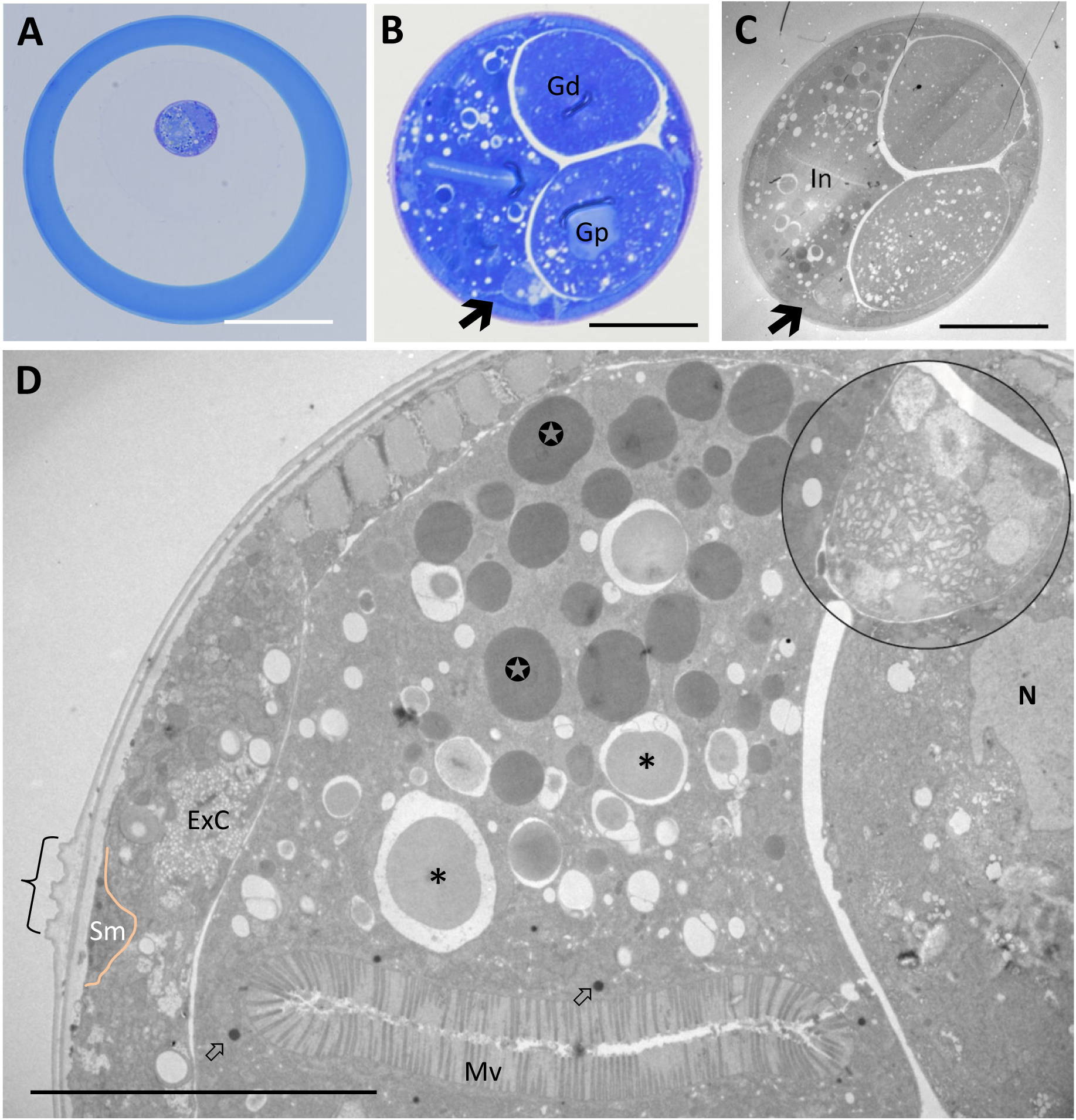
Low power cross section views of *C. elegans*. A and B, light micrographs of semi-thin sections stained with toluidine blue. C and D, electron micrographs of ultrathin sections stained with uranyl acetate and lead citrate. A: worm enclosed within a hollow fiber; B and C: consecutive sections of worm. Note that the ellipitical profile of the worm in C is due to section compression. (In=intestine; Gd= distal gonad; Gp= proximal goand; arrow = coelomocyte). D: an upper left quadrant of a cross-sectioned worm. The colored line marks the basal boundary of a seam cell. (* = gut granules; ⬀ = yolk granules; ✪= lysosomes; Mv= microvilli; { = alae; Sm= seam cell ; ExC= excretory canal; N: nucelus of proximal gonadal arm; Inset and 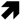 : coelomocyte). bar: A=100 μm, B and C = 20 μm, D= 10 μm

### Preservation of Internal Anatomy

Figure 4 depicts the preserved spatial arrangement of internal organs in *C. elegans*. Under physiological conditions, the worm’s elevated internal hydrostatic pressure functions as a “hydrostatic skeleton.” Disruption of this pressure—such as during dissection—often results in structural collapse or displacement. To prevent this, we applied intermittent microwave irradiation to enhance fixative penetration while the worm remained alive and intact (Table 1, step 1). This brief pre-fixation protocol effectively stabilized internal tissues prior to microdissection. Importantly, no extrusion of the intestine or gonads was observed during subsequent manipulation with micro scissors. The preserved anatomical relationships of major tissues and organs, as shown in Figure 4, confirm that microwave-assisted pre-fixation provided sufficient structural stabilization to maintain integrity following the release of hydrostatic pressure.

### Chemical Exchange Through EVOH Hollow Fiber

A central question in this study was whether EVOH hollow fiber might impede chemical exchange, thereby compromising infiltration and polymerization during sample preparation. To address this, we conducted a detailed ultrastructural examination of the *C. elegan*s midbody region. This anatomical zone was chosen for two primary reasons: (1) the midbody is more prone to processing artifacts than the head due to its fluid-filled pseudocoelom and elevated internal pressure, making it a more sensitive target for evaluation; and (2) all seven major physiological systems—integumentary, muscular, nervous, digestive, reproductive, excretory, and coelomocyte—are represented within this region (Fig. 4B–D). High-magnification TEM analysis, presented below, demonstrates the preservation and quality of sample preparation using the EVOH encapsulation method.

### Body Wall Ultrastructure

The body wall of *C. elegans* consists of three principal tissue layers: the cuticle, hypodermis, and obliquely striated musculature. As shown in Figures 4D and 5A, the cuticle exhibited successive tiers of organization and remained closely apposed to the underlying hypodermis. In nematodes, detachment of the cuticle is indicative of osmotic stress (Jones and Gwynn, 1991; Kirk et al., 2002). Thus, the observed integrity of the cuticle–hypodermis interface suggests that our sample preparation protocol did not induce osmotic shock. The musculature beneath the hypodermis was also well preserved. Muscle cells displayed two distinct domains: (i) the sarcomeric region, where the myofibrillar lattice, dense bodies, and M-lines were clearly defined and well organized; and (ii) the muscle belly, which contained ribosomes, endoplasmic reticulum, mitochondria, and nuclei—all exhibiting characteristic morphology and preservation. Notably, junctional complexes anchoring the body wall components were sharply delineated. These included fibrous organelles within the hypodermis (Fig. 5B), attachment plaques in muscle cells (Fig. 5B), and adherens junctions between seam cells and hypodermis (Fig. 5C). Together, these features underscore the structural fidelity of the body wall achieved through our optimized preparation protocol.

**Figure 5:**
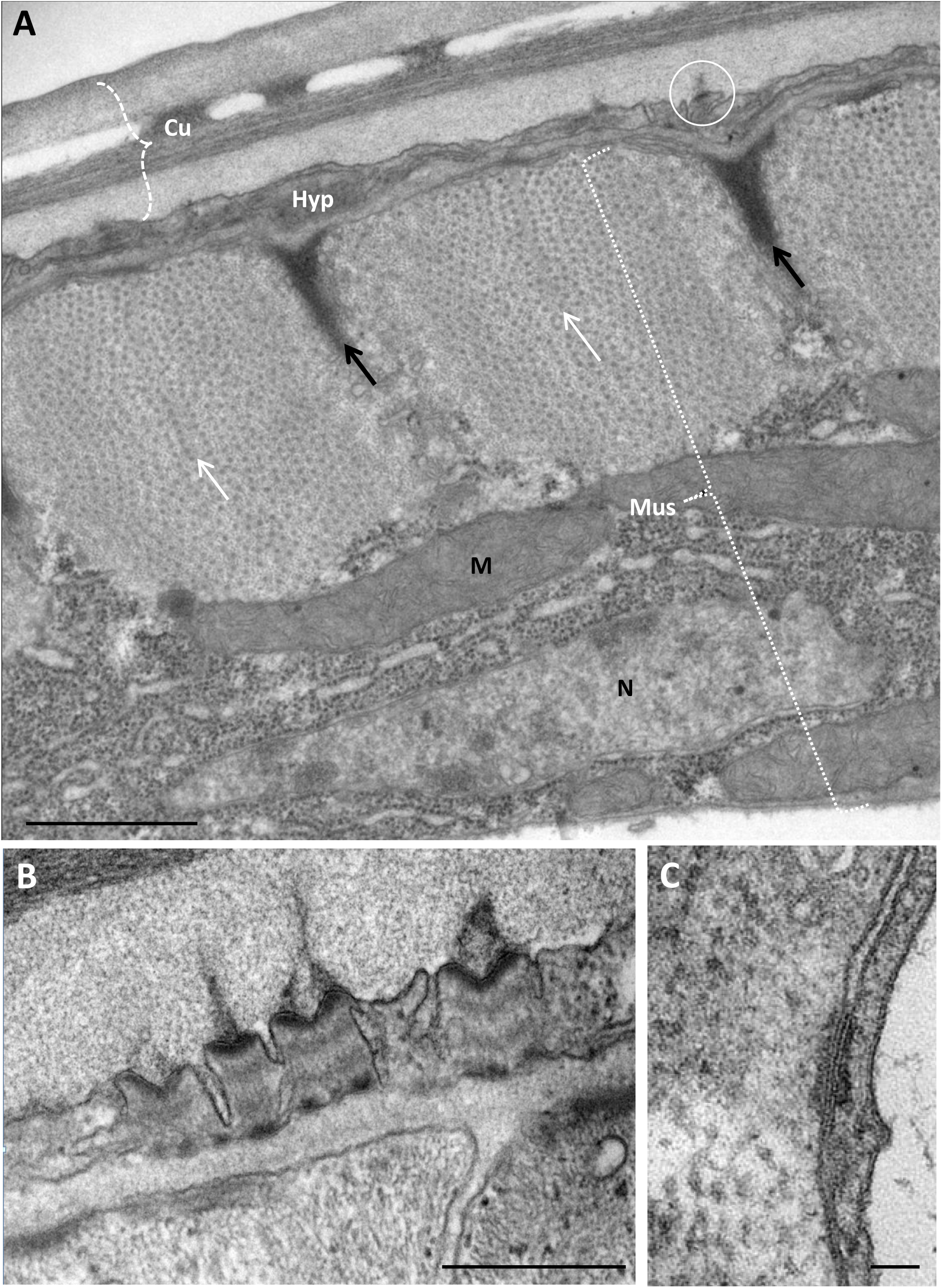
Body wall structures. A: The three layers of body wall are: cuticle (Cu), hypodermis (Hyp), muscle (Mus). ( N = nucleus; M = mitochondria; black arrow = dense body; white arrow = M line). B: fibrous organelle, (see also the encircled structure in A). C: adherens junction between seam cell and hypodermis. bar: A=500 nm, B=200 nm, C=20 nm.

### Membrane Organelles and Contacts

Our protocol yielded favorable preservation of intracellular membrane systems, a critical concern given the susceptibility of biological membranes to electromagnetic radiation and aldehyde-based fixation (Liburdy and Vanek, 1985; Murk et al., 2003). Figure 6 presents a panel of representative images illustrating a diverse array of membrane-bound structures and intercellular contacts. Observed features include multivesicular bodies, apical membrane stacks, and Ward bodies within hypodermal cells (Fig. 6A, 6B); yolk granules, secretory vesicles, endosomes, and apical junctions in intestinal cells (Fig. 6C, 6D); and the Golgi apparatus and gap junctions in excretory canal cells (Fig. 6E). These morphologies are consistent with conventional TEM descriptions and indicate that membrane integrity was well maintained throughout processing.

**Figure 6:**
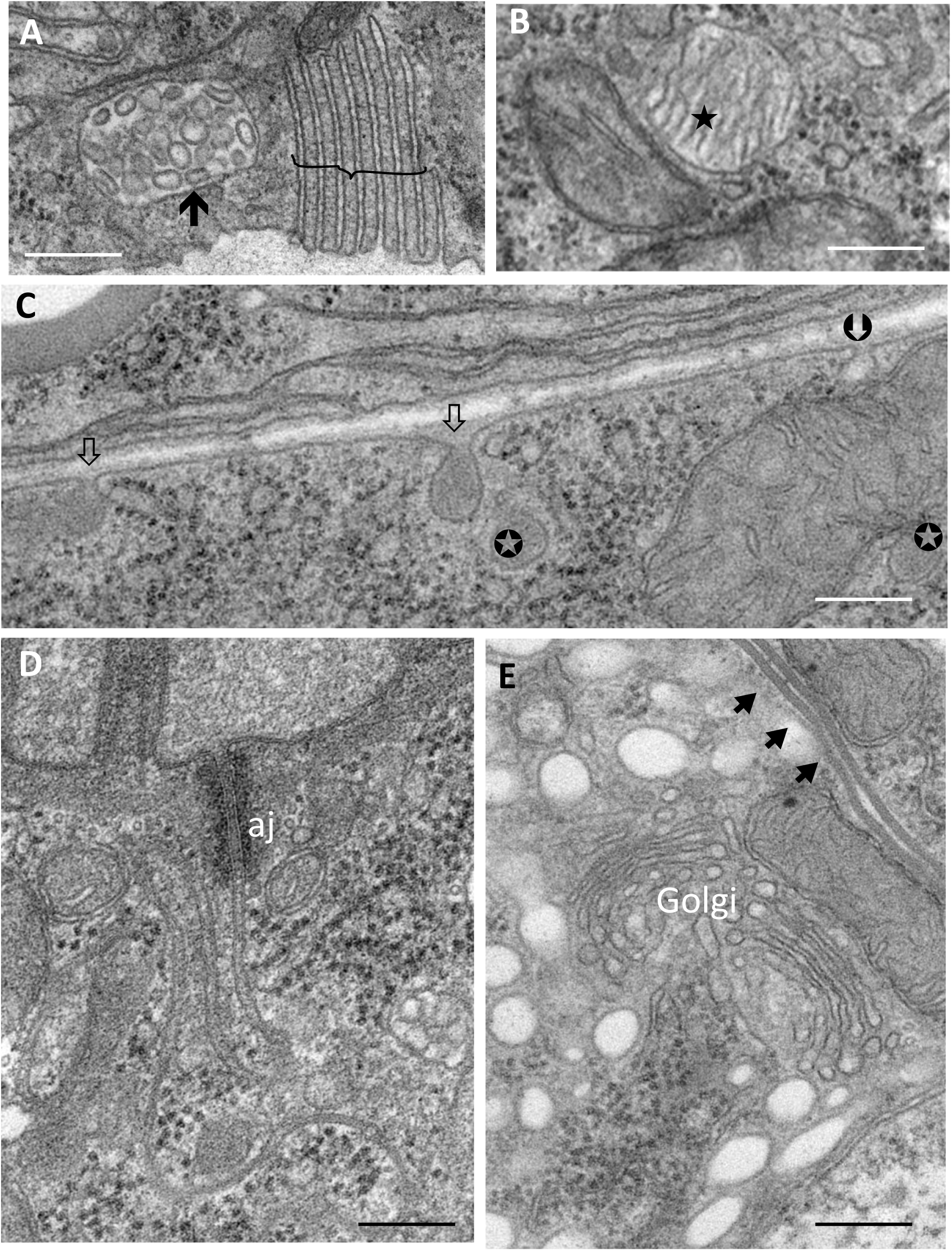
Membrane vesicles and appostions. Images of A and B are from hypodermal cells, C and D, intestinal cells, and E, excretory canal cell. (⬆ = multivescular body; } = apical membrane stacks; ★ = Ward body; ✪ = yolk granule; ⇓ = secretory yolk granules; 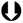 = endocytic pit; aj =apical junction; Golgi = Golgi apparatus; 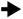 = gap junctions.) bar: 100 nm

### Neuronal Ultrastructure

Although not formally classified as part of the body wall, nervous tissue in the midbody region is closely associated with the hypodermis and merits inclusion here due to its prominence in electron microscopy studies of *C. elegans*. Figure 7 illustrates the high-quality preservation of neuronal ultrastructure achieved with the current protocol. Neurite profiles appeared smooth and turgid, and although mild undulations of the plasma membrane were observed—consistent with aldehyde fixation—these did not obscure surface invaginations (Fig. 7B). Presynaptic densities (open arrows, Fig. 7A, 7C, 7D) were readily identifiable, and both synaptic vesicles and dense core vesicles were distinguishable based on their morphology and spatial distribution. Sensory neurons were similarly well-preserved, as exemplified by the ALM touch receptor neuron embedded within the hypodermis beneath the cuticle. This neuron exhibited prominent large-diameter microtubules and a clearly defined extracellular mantle (Fig. 7E). To further demonstrate the efficacy of the infiltration protocol, Figure 8 presents tomographic data from a section that remained structurally intact under high-dose electron irradiation during a multi-exposure tilt series.

**Figure 7:**
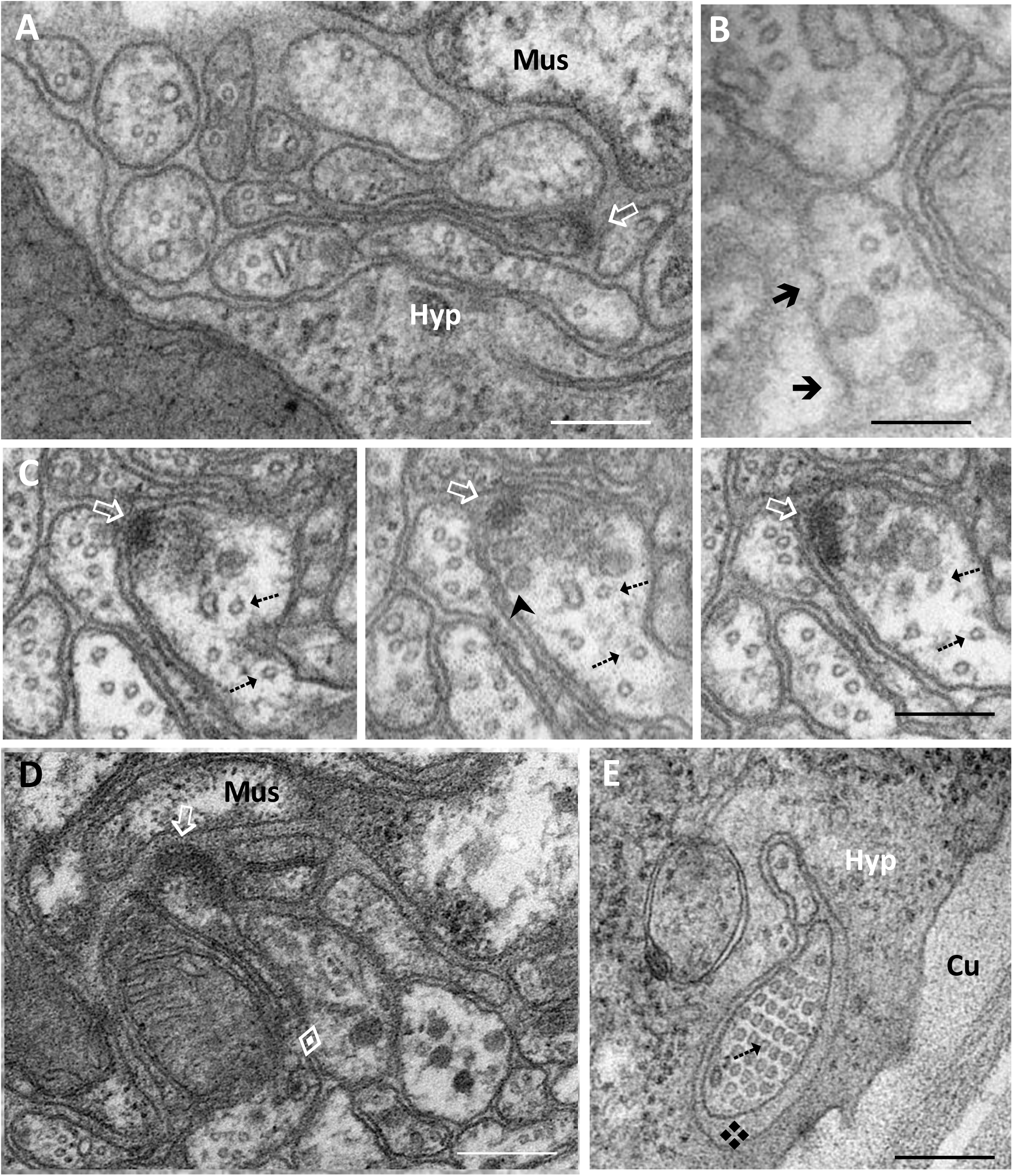
Ultrastructure of neurites. A: dorsal nerve cord; B-D: ventral nerve cord; E: sensory neuron ALM. (Hyp= hypodermis; Mus= muscle; 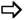 = presynaptic density; 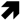 = surface invagination; 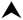 = synaptic vesicle; 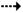 = microtubule; 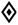 = dense core vesicle; 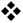 = mantle.) bar: 100 nm

**Figure 8:**
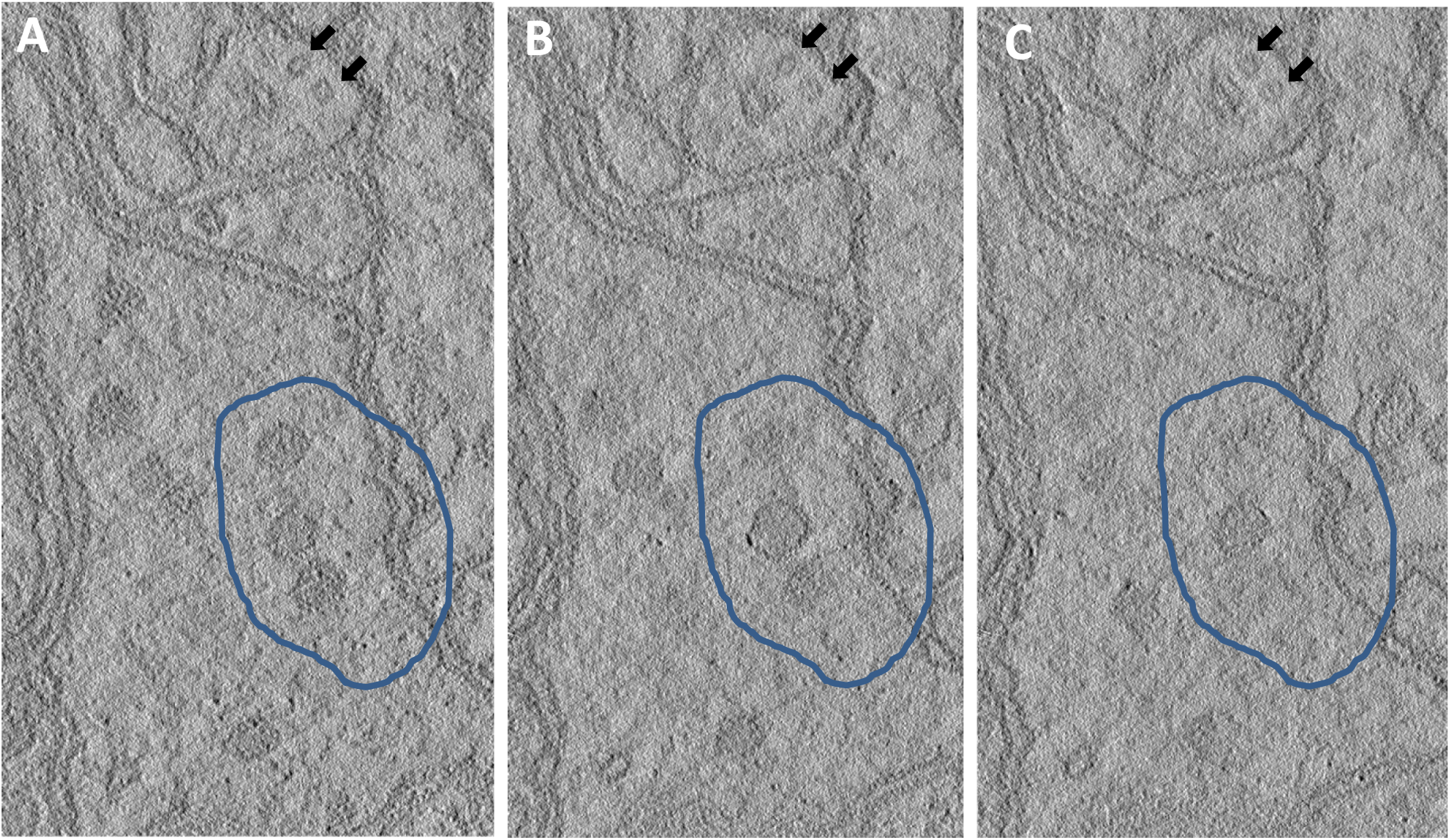
Electron tomographs of ventral nerve cord. Digital slices from the top, middle and lower end of an 80 nm section. In the highlighted area, spherical nature of neural vesicles are resolved. In contrast, the rod-shaped microtubules (⬅) is orthogonal to the plane of section.

## Discussion

### Advantages of EVOH over Cellulose-Based Capillary Hollow Fibers

The principle of encapsulating minute biological samples within capillary dialysis tubing was first introduced by Hohenberg et al. (1994) and later adopted by several researchers (Claeys et al., 2004; Matsko and Mueller, 2005; Muller-Reichert et al. 2003) in the context of high-pressure freezing of living specimens. In these earlier studies, the capillary tubes were cellulose-based, a material that tends to collapse upon drying, thereby limiting its use to the hydrated state.

In contrast, EVOH (ethylene vinyl alcohol copolymer) hollow fibers retain their lumen shape even after 10 years of dry storage. The persistent and well-defined lumen of EVOH fibers inspired the use of an insect pin for specimen handling, offering a practical and precise method for manipulation. Notably, encapsulation using EVOH fibers is not restricted to aqueous environments; it can also be performed with samples suspended in organic solvents. For example, we successfully encapsulated *C. elegans* specimens while they were soaked in a 1:1 mixture of resin and acetone. This remarkable versatility is likely due to EVOH’s unique polymer composition, which includes both hydrophilic and hydrophobic segments, allowing the fiber to maintain a distended, rounded contour under a wide range of processing conditions.

Another notable difference lies in the physical dimensions of the tubing: the EVOH fibers used in this study have a luminal diameter of 175 μm, compared to >200 μm in previous cellulose-based applications. Given that an adult *C. elegans* worm measures approximately 50 μm in diameter, the narrower EVOH tube offers a clear advantage by restricting the worm’s alignment to the longitudinal axis of the fiber, which is critical for achieving consistent orientation during sectioning.

The wall thickness and pore size—which correspond to the molecular weight cut-off (MWCO)—also differ significantly between the two materials. Cellulose tubing typically has a 15 μm wall thickness and a 5 kDa MWCO, whereas EVOH fibers feature a 25 μm wall thickness and a 30 kDa MWCO. Our results indicate that the thicker-walled EVOH fibers provide greater mechanical rigidity, while the larger pore size facilitates free exchange of all processing reagents, making EVOH hollow fibers highly suitable for resin-based sample preparation workflows.

### Comparison of Hollow Fiber with Gelatinous Media

One feature that distinguishes hollow fiber encapsulation from methods using gelatinous media is the pre-formed uniformity of the fiber. Hollow fibers offer precise and consistent dimensions, unlike hand-trimmed gelatinous blocks, which can vary in size and shape. This dimensional consistency facilitates accurate orientation of specimens. A key application that benefits from this property is cryosectioning of hydrated specimens, where freezing— rather than resin embedding—is used to harden the specimen for sectioning (Tokuyasu, 1973)

Another promising area of application is the encapsulation of cells and small colonies (data not shown), particularly when working with low cell numbers, such as post-sorting populations. In such cases, hollow fibers offer a distinct advantage: concentrated cell suspensions can be directly withdrawn into the fiber, bypassing the need for repeated centrifugation steps that often result in cell loss. This is especially relevant for rare cell populations, such as hematopoietic stem cells, where traditional TEM sample preparation methods require millions of cells and often fail to yield meaningful data.

## Conclusion

This article presents a streamlined procedure for the efficient encapsulation of *C. elegans* specimens, serving as a proof of concept for handling scarce or small biological samples in TEM analysis. Additionally, we describe several ancillary techniques that are simple yet effective for maintaining single-worm identity prior to ultramicrotomy. These include the use of a transparent silicone mold, wire loop, and micro-scissors. Collectively, these tools expand the existing methodological toolkit for *C. elegans* research.

## Acknowledgments

We thank Asahi Kasei Medical Trading (Taiwan) Co., Ltd. for providing the Kawasumi Laboratories dialyzer (model KF-201, approval number 15900BZZ01521) used in this study. We also thank Nancy Chandler for establishing the microwave protocols for the EM core of HSC, University of Utah. Finally, we acknowledge the support of the Cell Imaging Core at the University of Utah for the use of the ZEISS Axio Scan.Z1 and Dr. Michael Bridge for his assistance with image acquisition.

